# Phenotypic subtyping via contrastive learning

**DOI:** 10.1101/2023.01.05.522921

**Authors:** Aditya Gorla, Sriram Sankararaman, Esteban Burchard, Jonathan Flint, Noah Zaitlen, Elior Rahmani

**Affiliations:** Bioinformatics Interdepartmental Program, University of California, Los Angeles, Los Angeles, CA, USA; Department of Computer Science, University of California, Los Angeles, Los Angeles, CA, USA; Department of Computational Medicine, David Geffen School of Medicine, University of California, Los Angeles, Los Angeles, CA, USA; Department of Human Genetics, University of California, Los Angeles, Los Angeles, CA, USA; Department of Medicine, University of California, San Francisco, San Francisco, CA, USA; Department of Bioengineering and Therapeutic Sciences, University of California, San Francisco, San Francisco, CA, USA; Department of Psychiatry and Behavioral Sciences, Brain Research Institute, University of California, Los Angeles, Los Angeles, CA, USA; Department of Neurology, David Geffen School of Medicine, University of California, Los Angeles, Los Angeles, CA, USA

## Abstract

Defining and accounting for subphenotypic structure has the potential to increase statistical power and provide a deeper understanding of the heterogeneity in the molecular basis of complex disease. Existing phenotype subtyping methods primarily rely on clinically observed heterogeneity or metadata clustering. However, they generally tend to capture the dominant sources of variation in the data, which often originate from variation that is not descriptive of the mechanistic heterogeneity of the phenotype of interest; in fact, such dominant sources of variation, such as population structure or technical variation, are, in general, expected to be independent of subphenotypic structure. We instead aim to find a subspace with signal that is unique to a group of samples for which we believe that subphenotypic variation exists (e.g., cases of a disease). To that end, we introduce Phenotype Aware Components Analysis (PACA), a contrastive learning approach leveraging canonical correlation analysis to robustly capture weak sources of subphenotypic variation. In the context of disease, PACA learns a gradient of variation unique to cases in a given dataset, while leveraging control samples for accounting for variation and imbalances of biological and technical confounders between cases and controls. We evaluated PACA using an extensive simulation study, as well as on various subtyping tasks using genotypes, transcriptomics, and DNA methylation data. Our results provide multiple strong evidence that PACA allows us to robustly capture weak unknown variation of interest while being calibrated and well-powered, far superseding the performance of alternative methods. This renders PACA as a state-of-the-art tool for defining *de novo* subtypes that are more likely to reflect molecular heterogeneity, especially in challenging cases where the phenotypic heterogeneity may be masked by a myriad of strong unrelated effects in the data.

**Code Availability:** PACA is available as an open source R package on GitHub: https://github.com/Adigorla/PACA

## 1 Introduction

Heterogeneity in the underlying biological mechanisms of complex disease and traits has practical and clinical implications: undifferentiated cases of a disease may represent the action of a variety of underlying causal processes, each of which may have a different prognosis or respond to a different treatment. One classic example is the differentiation of diabetes mellitus into type 1 (insulin responsive) and type 2 (non-responsive) [1]. Other examples include the sub-classification of autoimmune thyroid disease into Hashimoto thyroiditis and Graves’ disease, the most common causes of hypothyroidism and hyperthyroidism, respectively [2], and the stratification of breast cancer patients based on estrogen receptor (ER) proteins on the surfaces of tumor cells (ER+/-), which is indicative of prognosis and response rates for drugs that interact with ER [3].

The term “phenotypic heterogeneity” is often used to describe observed variation of a given phenotype between individuals, irrespective of whether heterogeneity in the molecular background of the phenotype contributes to that variation. Here, we consider phenotypic heterogeneity in the sense of variability in the molecular basis of a phenotype. Such heterogeneity reflects possible variation or difference in the mechanism of the phenotype across individuals, which may be driven by genetics [4–10], environmental exposures and/or by the interaction thereof [10–18]. In fact, phenotypic heterogeneity is suspected yet unknown or not fully characterized for many conditions, including neurodegenerative diseases (e.g., Alzheimer’s disease [19] and dementia [20]), autoimmune diseases (e.g., rheumatoid arthritis and systemic lupus erythematosus [21]), and psychiatric disorders (e.g., major depressive disorder (MDD) [22, 23] and post-traumatic stress disorder [24]).

In the absence of the right subgrouping, phenotypic heterogeneity may compromise statistical analysis by leading to a substantial power loss and potentially low reproducibility rates in detecting and understanding the underlying mechanisms of heterogeneous phenotypes [10, 25–27]. Since the sub-classification or heterogeneity nature of the molecular background of phenotypes is typically unknown, it becomes a computational and statistical challenge to find surrogates to the subtypes.

Numerous studies have defined phenotypic subtypes based on histology [28, 29], genomics [30–32], electronic health records [33], or other phenotypically- or clinically-determined scores (e.g., [23, 34–36]). This approach occasionally reveals disease subgrouping that facilitates the detection of novel genetic signals. For example, using clinical and familial criteria, subgroups have often been proposed for lithium responsive bipolar disorder [34] and mood incongruent psychosis [35]; association studies with MDD indicate that its underlying genetic basis differs between those who have and those who have not experienced adversity [23]; and subtype characterization using gene expression signatures identified classes of tumors with distinct patterns of chromosomal alterations [37]. However, heterogeneity in the molecular basis of disease might not be observed phenotypically (e.g., both type 1 and type 2 diabetes share some similar symptoms), and in general, ad hoc clinically- or phenotypically-derived subgrouping is not guaranteed to capture variation in the underlying mechanisms.

Arguably, the most common systematic approach for defining *de novo* subtypes in a data-driven fashion with no prior information is data clustering and dimensionality reduction. This subtyping approach employs methods ranging from classical clustering and dimensionality reduction methods, such as K-means [31], hierarchical clustering [38, 39], and principal components analysis (PCA) [40–42], to more modern deep-learning based methods, including various types of autoencoders and other generative network models (see [43] for a review). These methods can possibly be used in combination with feature selection or feature weighting based on domain-specific prior knowledge [44, 45]. Importantly, all of these methods are essentially geared towards learning latent representation of samples based on the dominant sources of variation in a given dataset. This renders such methods ineffective in the typical case where phenotypic heterogeneity is not mirrored by the top axes of variation in the data. Particularly, the most dominant sources of variation in biological data most often reflect various population- or sample-related factors that can be independent of causal phenotypic variation, such as population structure [46, 47], cell-type composition [48], and technical variation [49, 50].

In some cases, the most dominant sources of variation in a dataset can lead to meaningful subtyping – perhaps most notably in genomic data from tumors, which include strong structure induced by the large number of mutations and genomic changes with cancer [52, 53]. However, the molecular signals underlying complex traits are most often expected to be weak (i.e., small effect sizes) and sparse; that is, the vast majority of the features available in a given dataset are not causal or statistically correlated with the phenotype of interest [54, 55]. Adjusting or conditioning the analysis on factors (i.e., as covariates) that are not expected to be statistically related to the subphenotypic signal can in principle alleviate this problem. Yet, in practice, unknown factors with unknown effects typically exist, and some of those factors may obscure the subphenotypic signal unless accounted for in the analysis. Therefore, whether conditioning on known covariates or not, a standard approach that relies on dominant variation in the data, in general, is not expected to allow us to robustly target phenotypic heterogeneity (or, more generally, a particular unknown weak variation of interest; Figure 1a-d).

**Figure 1:**
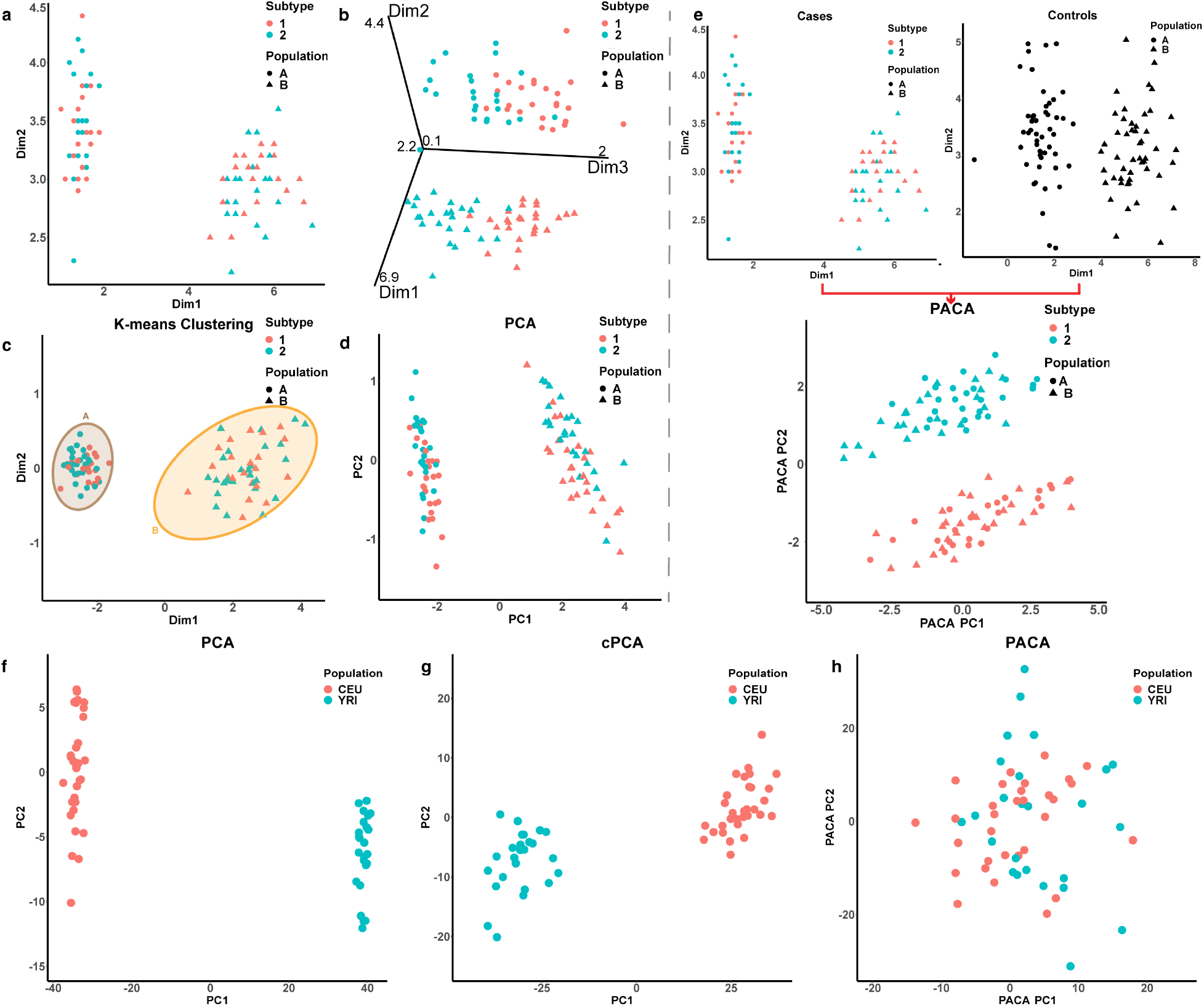
An illustration of learning weak sub-phenotypic signals. (a) Dimensions 1 and 2 of the data stratify samples by population (circles/triangles) rather than by subtype (red/green). (b) A third dimension of the data, which reflects a weaker signal, stratifies observations into the two subtypes. (c-d) A standard K-means clustering approach and dimensionality reduction using PCA surface population structure, the dominant source of variation in the data. (e) Contrasting cases with controls (which include no subtype signal) allows PACA to isolate cases-specific variation (i.e., subtypes), regardless of the rank of the signal. (f)-(h) Evaluation of the top two components of PCA, cPCA, and PACA under a scenario with no subphenotypic heterogeneity, in which samples from the HapMap data [51] were randomly assigned with a case/control status (58 YRI and 58 CEU individuals; based on a randomly selected set of 10,000 SNPs). Two outlier samples were excluded for visualization purposes).

Given the potential impact of identifying phenotypic heterogeneity on our ability to improve healthcare outcomes, an important question is thus whether we can systematically learn *de novo* phenotypic subtypes in cases where the phenotypic heterogeneity constitute only a small portion of the variation in the data and no prior information on the nature of the phenotypic heterogeneity is known.

### Related work

Targeting weak sources of variation in a dataset can in principle be achieved by conditioning on background data and introducing subspace-specific regularization [56]. More recently, a method following this approach, contrastive PCA (cPCA) [57], has been proposed and applied for the task of subtyping and sample subgrouping using protein and gene expression data. cPCA seeks to learn directions of variation that are enriched in a target dataset (e.g., cases) compared a background dataset (e.g., controls), which reflects a contrastive learning paradigm by which a weak form of supervision (target versus background labels) informs the unsupervised learning of patterns in the target data. In theory, the contrastive learning approach allows us to study subphenotypic variation in a group of cases by accounting for all variation that exists in controls, thus, dismissing the need for explicitly considering the (unknown) right set of covariates if taking a more straightforward approach of analyzing the cases alone.

The cPCA algorithm was demonstrated to successfully capture subgrouping information on several tasks [57], however, it is expected to be limited in scenarios of unsupervised detection of *de novo* subtypes where we do not have a prior knowledge about the right level of subspace regularization. Concretely, cPCA indicates for the user several possible levels of subspace regularization based on similarity of their induced subspaces in terms of their principle angles, yet, this may lead to tagging confounders. In the absence of a priori knowledge of subtypes, the analyst is likely to misspecify the regularization parameter, which may lead to falsely tagging arbitrary patterns in the target data as subphenotypic variation (Figure 1g). Critically, since biological confounders tend to replicate across independent datasets from the same population, tuning the regularization parameter via standard cross validation procedures is in general not possible.

Finally, deep learning methods for contrastive learning exist (e.g., contrastive variational autoencoders [58]), however, they require large sample sizes in order to learn models with a large number of parameters and address the sensitivity of deep learning to architecture and hyperparameter tuning. Given that large sample sizes are often not available in biological datasets, we do not consider such methods in our evaluation.

### Contributions

We propose Phenotype Aware Components Analysis (PACA), a method for robustly capturing weak variation of interest in high-dimensional data. PACA is a contrastive learning algorithm leveraging Canonical Correlation Analysis (CCA) to learn patterns in a given target dataset that cannot be found in background data. Given case-control data of any modality, PACA highlights the cases-specific dominant variation of the subspace that is not affected by control variation as a putative representation of phenotypic heterogeneity. By defining background variation as any biological or technical variation that is shared between cases and controls and is stronger than the cases-specific variation in the data, we cast PACA as an estimation algorithm, which subverts the need for a vague and hard-to-tune contrastive hyperparameter.

## 2 Methods

### 2.1 Preliminaries

#### Problem setup

We consider a low-rank model for high-dimensional data coming from a target population and a background population. Both populations share the same biological and non-biological sources of variation, with the exception of an additional source of variation that is unique to the target population and represents subphenotypic signal. Hereafter, we refer to the target and background groups as “cases” and “controls”, as a representation of the status of the samples in the data with respect to a condition of interest.

Let *X* ∈ ℝ^*m*×*n*_1_^ be a matrix of measurements of *m* features for *ni* cases, and let *Y* ∈ ℝ^*m*×*n*_0_^ be a matrix of measurements of the same *m* features for no control samples, such that *m* > max(*n*_0_,*n*_1_). We consider the following descriptive model:

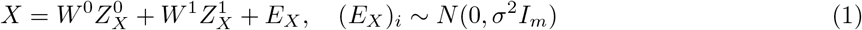

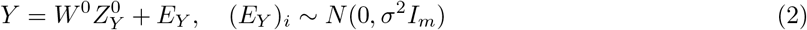

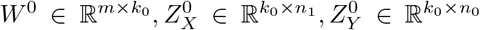 represent the directions (in the features space ℝ^*m*^) of a *k*_0_-dimensional low-rank signal (typically *k*_0_ << min(*n*_0_,*n*_1_)) and sample-specific structures for the samples in *X, Y*, respectively, by which the shared sources of variation across cases and controls are encoded. 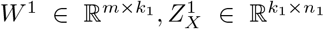 represent the directions of a *k*_1_-dimensional low-rank signal and sample-specific structure for the samples in *X*, respectively, by which the cases-specific sources of variation are encoded.

Since 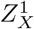 represents variation that cannot be attributed to any source of variation that exists in control samples it can be interpreted as variation due to heterogeneity among cases. In our context, we refer to this cases-specific variation as phenotypic heterogeneity or phenotypic subtypes; of note, while the latter may imply a categorical classification of samples into a constant number of groups, the former conceptually allows a more flexible characterization of the cases-specific variation by considering a spectrum to represent the phenotypic heterogeneity. Our goal is to learn 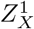 up to a linear transformation. We are particularly interested in a regime in which the cases-specific signals are weaker than (at least some of) the sources of variation that are shared across cases and controls (i.e., *W*^1^ spans less variation compared to *W*^0^).

#### Orthogonality assumption

If *W*_1_ is in the subspace spanned by *W*_0_ then all the structure in *X* and *Y* is spanned by the same subspace. This case corresponds to a scenario in which there is no cases-specific variability in the data, and although subphenotypic variation may exist, 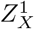 appears non-identifiable without additional supervision. Here, we make the assumption that *W*^0^ ⊥ *W*^1^. That is, we assume that the axes of common variation across cases and controls are orthogonal to the cases-specific variation. This assumption suggests that under the model in Eq. (1)–(2) a sensible approach for learning the cases-specific variation is to find sources of variation in *X* that do not exist in *Y*. Neglecting this assumption, while not allowing further supervision, may lead to under-correction for shared sources of variation across cases and controls. This, in turn, can lead to falsely tagging background variation as phenotypic heterogeneity (Figure 1f-h).

Notably, linear effects with the condition under study, in general, do not describe a dichotomous relation between features and the condition but rather a statistical one. Therefore, features with such linear effects are typically expected to vary in control samples too. Yet, this for itself does not nullify the orthogonality assumption, which does not concern feature-specific variation but rather reflects the assumption that control samples do not exhibit a systematic variation that is unique to cases. That being said, in practice, the orthogonality assumption may be violated since phenotypic heterogeneity may also be correlated with factors that exist in controls, such as population structure and demographics. As we later show empirically, violation of the orthogonality assumption will lead to a decrease in sensitivity to capture cases-specific variation. However, working under this assumption has the benefit of avoiding under-correction for the shared variation across cases and controls.

#### Contrastive PCA is limited under the orthogonality assumption

Using the same notations as above, given a parameter *α* > 0, cPCA finds the top contrastive principal component (PC) by solving [57]:

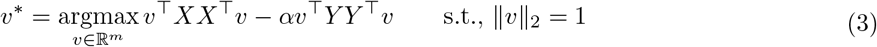

In words, cPCA aims at finding the dominant axis of variation in *X* while controlling at a level associated with α for the variation this axis describes in *Y*. This framework allows us in principle to learn the cases-specific variation 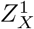 described in Eq. (1)–(2). Particularly, setting *α* = ∞ allows cPCA to incorporate the orthogonality assumption and account for all shared sources of variation across *X* and *Y*. A solution to Eq (3) while setting *α* = ∞ requires that ||*Y*^τ^*v*\*|| = 0 in order to avoid a negative infinity value for the objective. Therefore, *v*\* must be orthogonal to the subspace spanned by the columns of *Y*, i.e., *v*\* ∈ null(*Y*): = {*v*|*Y*^τ^*v* = 0_*m*_}, where 0_*m*_ is an *m*-length vector of zeros. Effectively, since *X, Y* are full-rank matrices due to the i.i.d. components of variation in Eq. (1)–(2), *v*\* must be orthogonal to an *n*_0_-dimensional subspace defined by to all samples in *Y*.

If *n*_0_ ≥ *m* then *v*\* must be orthogonal to the entire *m*-dimensional features space, hence we get *v*\* = 0_*m*_ irrespective of the rank of *X*. Here, we are interested in the case where max(*n*_0_, *n*_1_) < *m*, therefore *v*\* is nontrivial, however, following the rank-nullity theorem, *v*\* is restricted to an *m*–*n*_0_ dimensional subspace of ℝ^*m*^. Clearly, as *n*_0_/*m* approaches 1, cPCA will not allow us to capture the direction that spans the subphenotypic signals. Intuitively, this approach is suboptimal since it conditions on the entire n_0_-dimensional subspace in *Y* rather than on the lower-dimensional subspace that captures only the low-rank signals in the data. This suggests that a better approach is to condition only on sources of variation that exist in both *X, Y*, which are expected to be of lower dimension compared with the dimension of the observed data. This is exactly the key idea behind PACA.

### 2.2 PACA: Phenotype-Aware Component Analysis

#### Capturing subphenotypic variation using PACA

Using a singular value decomposition we can consider an alternative formulation for the model in (1)-(2):

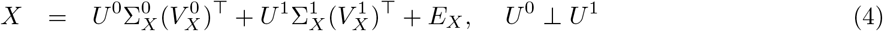

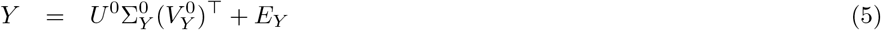

where each of 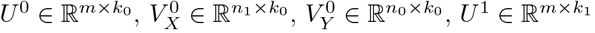, and 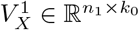 forms an orthonormal basis. Under this presentation, we are interested in learning 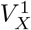, a linearly-transformed surrogate for 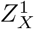 in Eq. (1). Given the directions of the shared sources of variation *U*^0^, note that

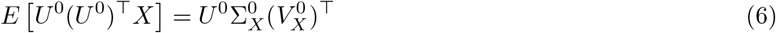

which established a way to remove the expected effect it induces on *X* (i.e., the signals in *X* that are coming from sources of variation that exist in controls). Our PACA algorithm therefore estimates and removes the effects of *U*^0^ on *X*, followed by PCA for capturing 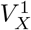. Our main focus is thus estimating *U*^0^.

Since *U*^0^ is found in both *X, Y*, we can estimate it by employing a canonical correlation analysis (CCA) [59]. Importantly, unlike in a typical application of CCA, where we wish to find linear transformations of the features that yield the highest correlation between the samples in two datasets (i.e., vectors in the samples space), here, we seek linear transformations of the samples that provide vectors in the features space ℝ^*m*^. Specifically, we find the first pair of canonical variables by solving:

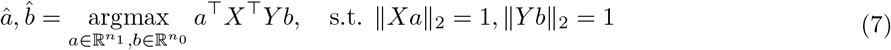

Setting 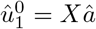 yields the representation in *X* of the strongest direction of shared variation across *X* and *Y*. Let *S_XY_* be the empirical sample cross-covariance of the matrices *X* and *Y*, the solution for 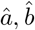 is known to be given by the eigenvector that corresponds to the top eigenvalue of 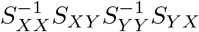 and the eigenvector that corresponds to the top eigenvalue of 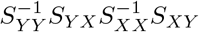, respectively [59]. This procedure can be repeated iteratively by restricting the vectors 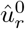 to be orthogonal to the previous canonical variables 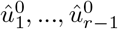 (and similarly for the variables of *Y*). Eventually, the collective of these vectors 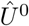 can be used for removing the shared sources of variation from *X* following Eq (6); see a summary of PACA in Algorithm 1. As we discuss next, choosing the number of components *r* to be used can be informed by the structure induced in *X* and *Y* by the shared variation.

##### Algorithm 1 Phenotype Aware Components Analysis (PACA)

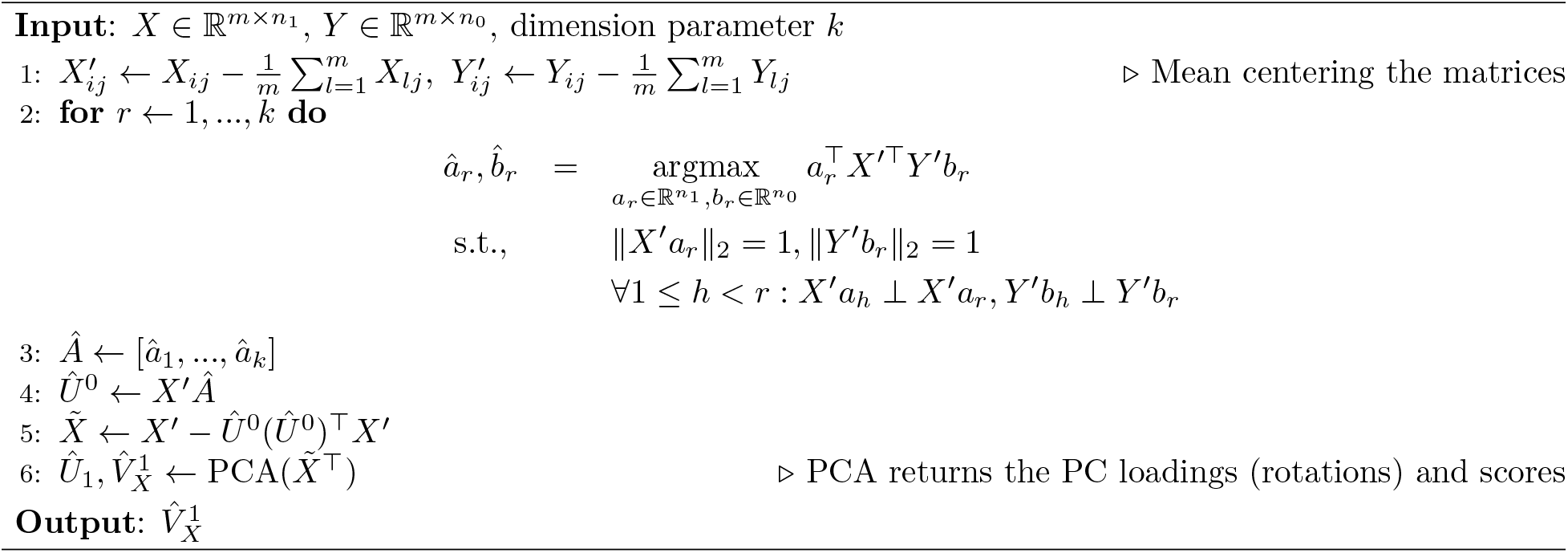

We note that the CCA step in PACA imposes the limitation *m* ≥ max(*n*_0_,*n*_1_). In practice, there may be datasets in which this condition does not hold. In such cases, we can in principle apply PACA on a subset of the samples, however, this is clearly sub-optimal to exclude data. Instead, we can use a randomized version of PACA that can operate under the setup *m* < max(*n*_0_, *n*_1_) by using multiple random subsets of the data for estimating the full sample covariance matrix of *X* (while accounting for the shared sources of variation between *X* and *Y*). See Supplementary Methods and Supplementary Algorithm 2 for details.

#### Estimating the dimension of the shared variation

In typical settings, the parameter k0 of the dimension of *U*^0^ is unknown. We may estimate it by setting an estimate 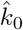 to be the largest value r that reveals a significant correlation between the correlated subspace of *X* and *Y* induced by 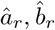; this can be achieved, for example using Bartlett’s Chi-squared test or Rao’s approximate F statistic [60]. However, in practice, we found that in the data we analyzed *k*_0_ seems to be very large (typically a few hundreds). Hence, this approach can lead to over-regularization and reduction in statistical power due to the unnecessary removal of a large number of axes of shared variation. In order to see why, recall that our goal is to be able to systematically target variation of interest that is typically weaker than the dominant sources of variation in the data. While standard dimensionality reduction tools cannot achieve this when applied to the original data, we expected them to be effective in revealing the target variation if applied to a residualized (adjusted) version of the data in which all sources of variation that are stronger than the target variation are removed. In other words, there is no need to estimate the true k_0_ and remove a structure of this dimension from the data prior to applying dimensionality reduction. Instead, we wish to find what is the minimal number of axes of variation k that we need to remove in order to reveal variation that is unique to cases. Below, we provide a brief description of the algorithm; see Algorithm 2 for complete details.

Given all shared axes of variation learned from the data using the procedure in Eq. (7), we learn *k* ∈ {1,…, min(*n*_0_,*n*_1_)} using binary search as follows. At each given candidate *k*, we evaluate the variance of the top PC we calculate from the residualized *X* accounting for the top *k* shared axes of variation; this same PC, which may reflect cases-specific variation, is also used for evaluating the variance it can explain in the residualized *Y* matrix. Comparing these variances to “null” variances obtained while permuting the loadings of the PC allows us to call whether accounting for *k* axes of shared direction is sufficient to detect structure that is unique to *X*. This evaluation can lead to one of four scenarios, based on which we decide on the next partition of the binary search: there can be (i) significant variation in cases but not in controls, (ii) significant variation in cases and controls, (iii) no variation in cases or controls, or (iv) significant variation in controls but not in cases.

Scenarios (i) and (iii) lead us to consider lower values of *k* due to possible over correction. The former means we revealed cases-specific variation, yet, we may be able to refine the signal if we can identify it using lower *k*, and the latter indicates that all the structure in the data has been removed. Scenario (iv) indicates a violation of our model assumption, which leads to termination, and lastly, scenario (ii) indicates residual shared variation, which suggests increasing *k*. Empirically, we found that considering a threshold *γ* on the maximum ratio between the variance of *X*’s PC and the variance it explain in *Y* under scenario (ii) is more stable and improves performance. Specifically, if the ratio between the variances of *X, Y* is greater than a predefined threshold then we handle this case similarly to case (i), and otherwise, we assume under-correction and consider a larger *k*. Throughout our analysis, unless stated otherwise, we used *γ* = 10.

### 2.3 Evaluation metrics and Datasets

See Supplementary Methods for complete details about datasets, data simulation, evaluation metrics, as well as complete technical details about the data analysis and implementation of PACA.

#### Algorithm 2 PACA: selecting *k* (the dimension of shared variation to remove)

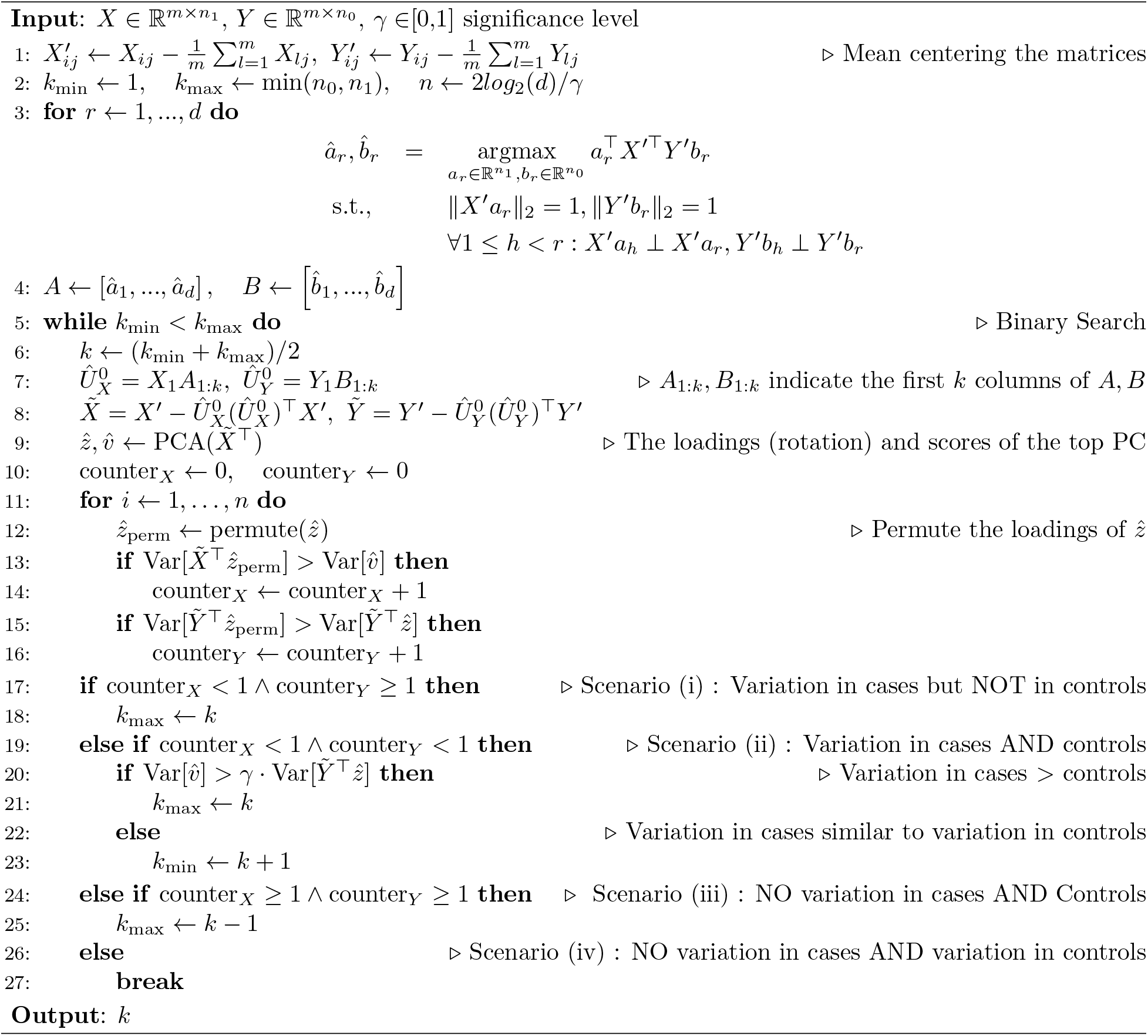

## 3 Results

### 3.1 PACA is calibrated under the null

A desired property of a method for capturing phenotypic heterogeneity is being calibrated under the null: given data with no phenotypic heterogeneity (i.e., the null case) and a prespecified confidence level. The method should be able to determine that no subphenotypic variation exists while controlling for the target type I error rate that corresponds to the prespecified confidence level. In order to verify that PACA is calibrated under the null, we applied it to data with no phenotypic heterogeneity by randomly sampling two groups of healthy individuals we labeled as “cases” and “controls” from each of the UK Biobank and the PsychENCODE datasets [61, 62](Supplementary Methods). We compared PACA with cPCA [57] and standard PCA, and using a permutation testing scheme we evaluated whether the structure found by each method represents phenotypic variation (Supplementary Methods). Empirically, we found that requiring cPCA to automatically select a single regularization parameter across a range of possible default values always favors no regularization (i.e., equivalent to PCA). We therefore set cPCA with a regularization parameter *α* = ∞, which we expected will favor the performance of cPCA in the calibration analysis. We found the PACA is calibrated and admits the lowest error rates among the different methods; for example, in the experiment using the PsychENCODE data PACA is calibrated with <5% error at a target type I error rate of 5%, yet cPCA and PCA demonstrate approximately 7% and 13% type I error, respectively (Supplementary Figure S1).

### 3.2 PACA is well-powered to capture phenotypic subtypes

We next evaluated PACA under the scenario where phenotypic heterogeneity does exist. To that end, we simulated high-dimensional data representing cases and controls. Specifically, every sample was generated by combining sample-specific structure with axes of variation that were shared across all samples. Then, we randomly split cases into two subtypes and added an additional weak and sparse cases-specific signal that differentiate between the two subtypes (Supplementary Methods).

In order to establish baseline performance we also considered standard PCA and cPCA. Since the latter depends on a regularization hyperparameter *α*, which is generally unknown, we considered two approaches. First, we applied cPCA while setting *α* = ∞, which allowed us to guarantee that cPCA learns variation that is strictly unique to cases. Second, we allowed cPCA to automatically recommend ten values of *α* based on the authors’ suggested algorithm for identifying subspaces that are maximally distinct from each other based the principle angles [57]. Then, we set the final *α* to the value that resulted in the best performance. This procedure represents a hypothetical scenario in which the subtype information is known and can be used for selecting the best *α*. This evaluation therefore reflects an over-optimistic evaluation of the performance of cPCA in practice. As before, setting cPCA to automatically select a single value of α always resulted in no regularization (i.e., *α* = 0, which is equivalent to a standard PCA).

Our results show that except for the case of very strong signals PACA outperforms cPCA in all settings, in spite of our over-optimistic evaluation of cPCA (Figure 2). Particularly, we observe that unlike cPCA, PACA is well-powered even in cases of very sparse signals (Figure 2). We further examined the performance of the different methods under a regime where the orthogonality assumption is violated and the subtype information is correlated with the shared sources of variation between cases and controls (Supplementary Methods). As expected, we notice a substantial drop in power for all methods when the orthogonality assumption is violated (Supplementary Figure S2). Notably, PACA appears to be the most robust method under this regime in cases of weak to moderate correlation between the subtype classification and the shared sources of variation (Supplementary Figure S2b). In cases of severe violation of the orthogonality assumption, the power of all methods drop considerably, effectively not allowing to detect subtypes (Supplementary Figure S2c,d)

**Figure 2:**
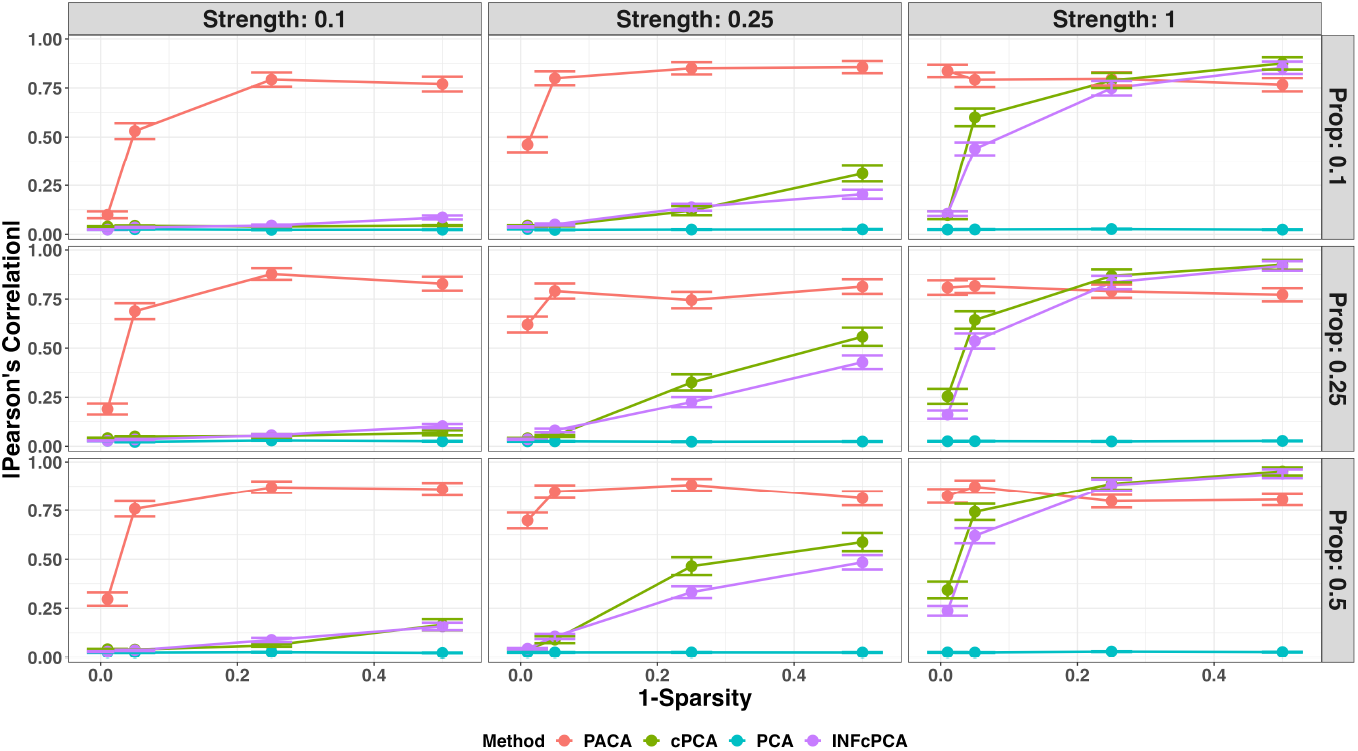
Evaluation of power to capture subphenotypic signal under simulations in the presence of two subtypes (*n* = 2000, *m* = 2000, *k*_0_ = 400). Performance was evaluated for PCA, PACA, cPCA with the best performing regularization parameter *α* among a range of values suggested by cPCA, and cPCA while setting the regularization parameter to *α* = ∞ (INFcPCA). Presented is the linear correlation of the top component of each method with the simulated subtype as a function of the signal sparsity (lower values mean more sparse), across a range of signal strengths (columns) and levels of imbalance (Prop) between the prevalence of the two subtypes (rows). The performance of every combination of parameters was averaged across 100 simulated datasets and vertical bars represent standard errors.

Finally, in order to gain more insight into the superior performance of PACA compared with cPCA, we compared the simulated dimension of shared variation between cases and controls (*k*_0_) to the dimension of shared variation that was removed by PACA. We found that in cases of weak subphenotypic signals PACA tends to adjust the data for a number of axes of shared variation that approximately matches the true simulated dimension of shared variation (Supplementary Figure S3). As the strength of the subphenotypic signal increases, a larger subspace of the space that defines the shared sources of variation is expected to include signals that are weaker than the subphenotypic signal. PACA leverages this insight and estimates the “effective” (i.e., minimal) dimension of shared variation that needs to be removed in order to detect subphenotypic signal (i.e., rather than removing all the shared variation; Algorithm 2); indeed, our analysis confirms that PACA adjusts for less axes of shared variation as the signal strength and density increase (Supplementary Figure S3).

### 3.3 Stratification of DNA methylation samples from admixed populations

The most dominant source of variation in DNA methylation from heterogeneous tissues such as blood is known to be cell-type composition [48], however it has been previously reported that a large number of methylation CpGs are strongly correlated with population structure [47, 63]. We therefore asked whether PACA can capture population stratification from heterogeneous methylation samples that were previously collected from individuals of Mexican and Puerto Rican descent (n=481) [48]. To that end, we applied PACA and cPCA to this target dataset while contrasting is with a background dataset of whole-blood methylation samples collected from European individuals (n=651) [64]. As expected, PACA achieved the best performance at capturing the target stratification in the data (Figure 3). On the other hand, cPCA could not capture the target variation either when considering the best performing regularization among a range of possible values of *α* suggested by cPCA (Figure 3c) or by setting *α* = ∞ (Figure 3d).

**Figure 3:**
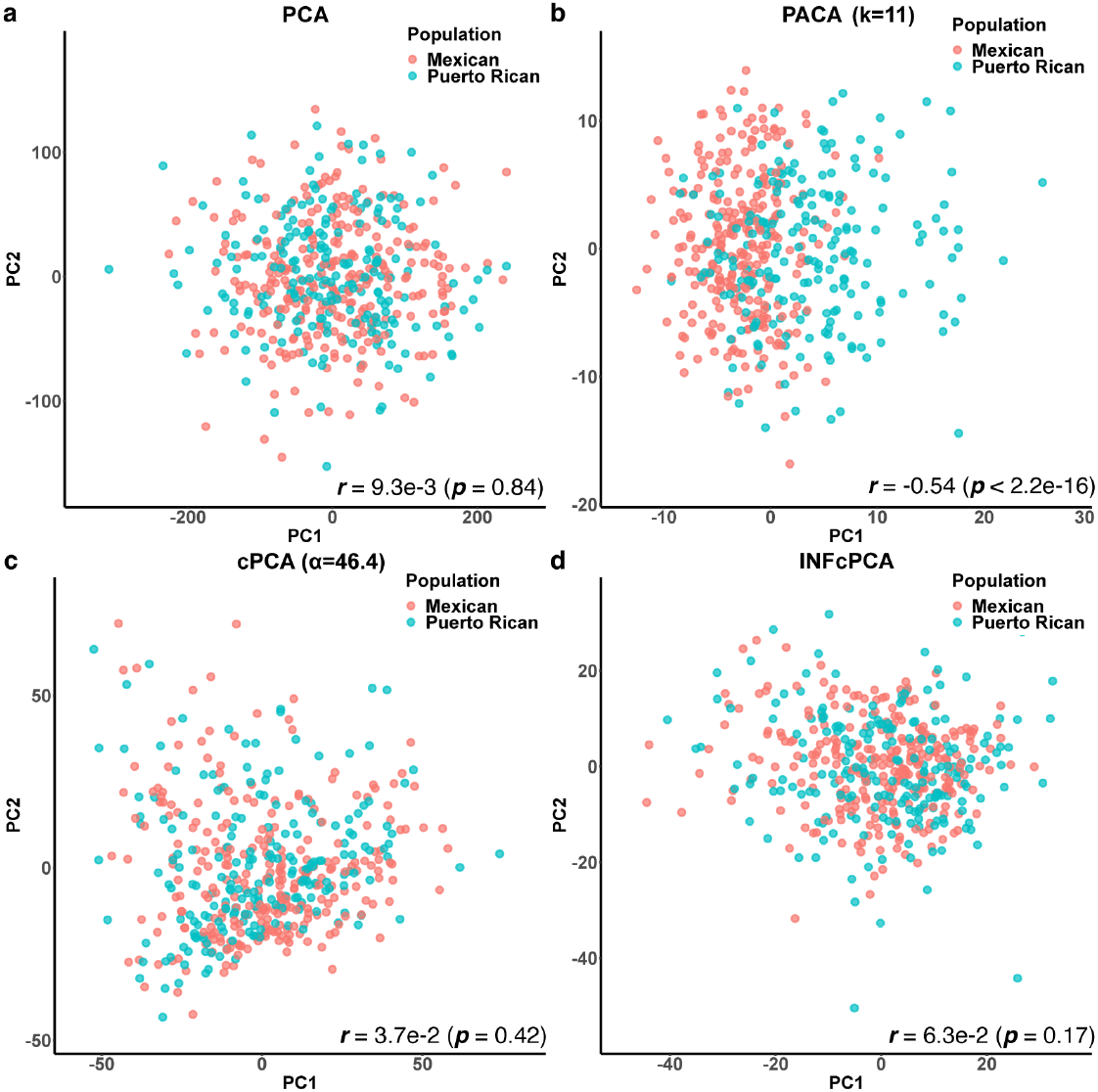
Capturing population heterogeneity in whole-blood DNA methylation data from admixed individuals. Presented are the first two components (denoted PC1 and PC2) found by (a) a standard PCA, (b) PACA, which accounted for an automatically estimated 11-dimensional structure of shared variation, (c) cPCA with the best performing regularization level among a range of values suggested by cPCA, and (d) cPCA with the regularization level set to *α* = ∞.

### 3.4 Identifying subphenotypic signal that reflects genetic heterogeneity under violation of the orthogonality assumption

Our simulation study suggests that PACA is more powerful than the alternatives in cases where the orthogonality assumption is not met, as long as the correlation between the subphenotypic signal and the shared sources of variation is weak or moderate (Supplementary Figure S2). In order to evaluate this result in real data, we applied PACA to a mixture of gene expression profiles from the PsychENCODE data [61, 62]. We pooled together schizophrenia (n=472) and bipolar (n=172) cases, the two most prevalent disorders in the data, as the “cases” group, thus, emulating a single disorder with two subtypes. For the background group, we considered all controls in the data (n=644). Prior to the analysis, we confirmed that the orthogonality assumption is violated: the simulated subtype was indeed correlated with the RNA sequencing technique used (library preparation) for the sample (poly(A) enrichment or ribosomal RNA depletion; r=0.12, p-value=0.003).

A simple application of PCA showed that the first PC of the cases group is correlated with the subtype assignment (r=-0.26, p-value=1.5e-11), however, this signal was driven by the correlation of the PC with the library preparation (r=0.70, p-value<2.2e-16). In fact, we found that the first two PCs of the data perfectly separates the samples by the library preparation type (Supplementary Figure S4). We therefore next asked whether applying PACA and cPCA using the controls group as a background data can capture subphenotypic signal that cannot be explained by the first two PCs of the data. While the first contrastive PC of cPCA did not capture substantial subphenotypic signal in that case (p-value=0.056 for the α resulting in highest correlation with the subtype; linear regression), the top PACA component of PACA did identify subtype signal when accounting for the first two PCs of the data (p-value= 2.6e-07; linear regression).

One way to evaluate whether the subphenotypic signal captured by PACA is meaningful (rather than merely reflecting other confounders) is by asking whether the subphenotypic signal captured by PACA is associated with genetic heterogeneity. A correspondence between the PACA component and changes in allele frequencies will provide strong indication for a real signal. In order to answer this question, we applied Subtest [65], a statistical method for the identification of genetic heterogeneity within phenotypically-defined subgroups (Supplementary Methods). Specifically, we applied Subtest on the genotypes of the PsychENCODE samples and provided it with the putative subphenotypic signal we learned using PACA. Indeed, Subtest confirmed that PACA captured an axis of subphenotypic variation that is indicative of differential genetics architecture among the cases (p-value <1.4e-6).

### 3.5 Capturing phenotypic heterogeneity in genetic data

Common genetic variation and specifically SNPs are notorious for their small effect sizes in complex disease [66, 67]. This makes the task of finding subtypes solely based on such genetic data particularly hard. We therefore evaluated the potential utility of PACA in identifying phenotypic heterogeneity from SNPs. To that end, we mixed genotype arrays of coronary artery disease (CAD) cases (n=2,500) and rheumatoid arthritis (RA) cases (n=2,500), which we pooled together from the UK Biobank data [68] into one group. Owing to the large number of genetic variants in typical genotype data, we worked under a relaxed assumption that prior work identified large sets of genetic variants that are likely to be associated with the phenotypic subtypes. Specifically, in our analysis, we considered the combination of a set of 6,000 random SNPs, the top 2,000 most associated SNPs with CAD, and the top 2,000 most associated SNPs with RA based on standard case-control GWAS (a total of 10,000 SNPs; Supplementary Methods). We applied PACA to this mixed group of cases while contrasting it with a group of healthy control individuals from the UK Biobank (n=5,000), and we found that the top PACA component is enriched for correlation with CAD- and RA-associated SNPs that represent a subphenotypic genetic heterogeneity in this case (Supplementary Figure S5a).

Finally, we applied Subtest to the data, which further confirmed that the top PACA component defines a spectrum of phenotypic heterogeneity that presents differential genetic architecture between the CAD and RA cases (p-value <4.201e-40). In total, we identify 211 SNPs (Subtest cFDR < 0.0434; Supplementary Methods) as likely to contribute to the difference in genetic basis of CAD and RA along the top PACA component (Supplementary Figure S5b).

## 4 Discussion

PACA is a contrastive learning algorithm for targeting weak sources of variation of interest from target data by leveraging background data that includes the same sources of variation that exist in the target data except for the variation of interest which is specific to the target data. Through an extensive simulation study and analysis of three genomic modalities, we demonstrated that PACA is calibrated and well-powered for identifying phenotypic heterogeneity. The power and robustness of our approach stems from the strategy of accounting for shared sources of variation across the target and background data. In particular, PACA aims at adjusting for the minimal number of shared axes of variation in the data that it is sufficient to remove in order to reveal variation that is specific to the target data. This strategy allows PACA to avoid falsely tagging confounders as subphenotypic signals and it subverts the need for a contrastive hyperparameter.

The robustness and calibration of PACA does come at a cost: our assumption that the subphenotypic signal is orthogonal to the confounders in the data may be violated in reality. A violation may occur either due to the nature of the subphenotypic signal or due to study-specific artefacts, such as case/control misclassification (i.e., noisy/incorrect labeling of controls as cases or vice versa). Not meeting this assumption may lead to power loss, however, we observe that PACA still supersedes the alternatives in scenarios of weak to moderate violation of the orthogonality assumption and comparable in cases of severe violation (Supplementary Figure S2). Furthermore, since PACA learns the directions of the shared sources of variation across *X, Y*, it can in principle be robust to imbalances in confounders between *X* and *Y*. However, much like any unsupervised method, PACA is expected to be sensitive to scenarios in which there are severe confounder imbalances or in cases where there are confounders that affect only *X* and not *Y*. If both *X* and *Y* were collected as part of the same study and under a proper randomization, such confounders in principle should not affect the data. Yet, in the event that such severe confounders are present in the data, they should be properly addressed. For example, in the case of multi-site data, if certain sites collected only cases or only controls then one should consider excluding the data from those sites prior to applying PACA. Otherwise, confounding effects due to site-specific variation will affect the ability of PACA to accurately capture the shared variation between cases and controls and the true cases-specific subphenotypic variation.

Finally, a straightforward implementation of the CCA backbone used by PACA restricts the number of samples to be lower than the number of features. That is, PACA is restricted to a “large-*p* small-*n*” regime (i.e., the number of features, p, is larger than the number of samples *n*). Yet, a desired property of a computational method is scalability and applicability to large sample sizes. In order to address this technical limitation of CCA, we developed a randomized version of PACA (rPACA), which allows us to apply PACA in regimes where *p* << *n* (Supplementary Methods).

## Acknowledgements

This research was conducted using the UK Biobank Resource under application 33127. We thank the participants of UK Biobank for making this work possible. This work was funded by the National Institute of Mental Health grant R01MH122569.

## Supplementary Materials for

### Part I

#### Supplementary Figures

**Figure S1:**
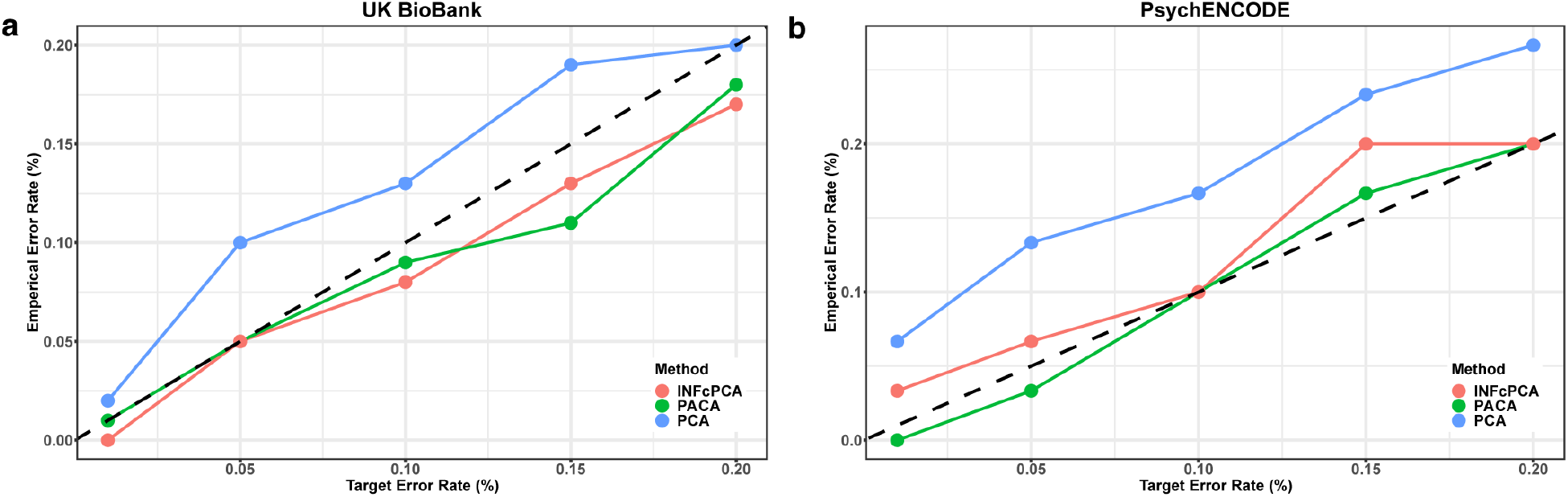
Evaluation of calibration under the null case of no subphenotypic variation. (a) The type I error calibration of PACA, cPCA (*α* = ∞; INFcPCA) and PCA on genotype data from non-diseased unrelated white British individuals from the UKBiobank (N=2,000 and a randomly selected subset of 10,000 SNPs) over 100 simulations. (b) The type I error calibration on gene expression data on control samples from PsychENCODE (N=1,000 and 25,774 genes) over 30 simulations. The X-axes represent the target type-I error and the Y-axes correspond to the observed type-I error at the target error rate. The dotted lines represent an ideal type-I calibration curve.

**Figure S2:**
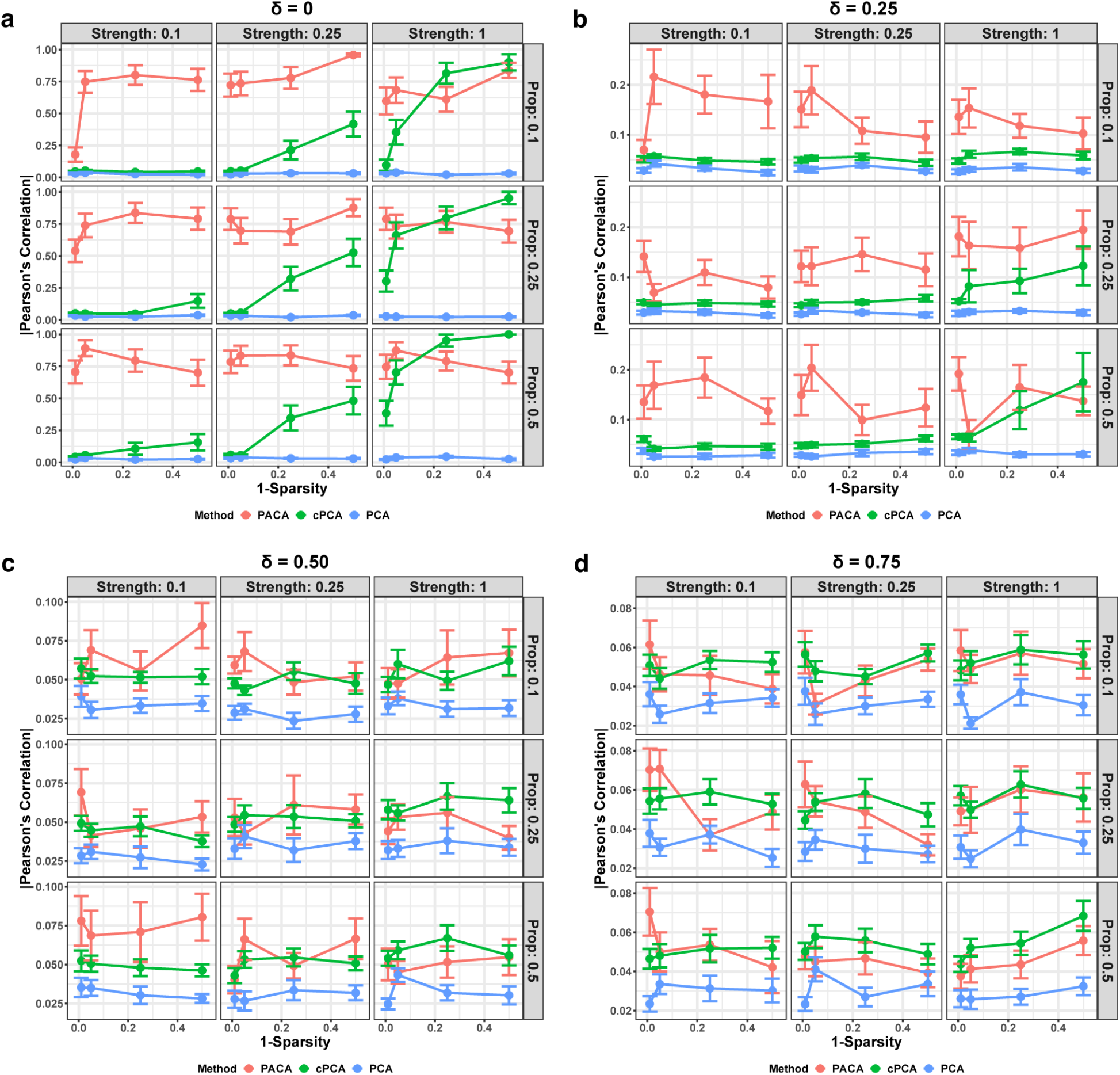
Evaluation of the power of different methods to capture subphenotypic signal under different levels of violation of the assumption of orthogonality between the cases-specific variability and the shared variability. For each of several levels of violation of the orthogonality assumption (denoted by *δ*), presented is the correlation of the outcome of the different methods with the subtype signal as a function of the signal strength, case subtype proportion (Prop), and signal sparsity (Supplementary Methods). *δ* corresponds to the ratio between the subtype variability in the controls and the subtype variability in the cases. Particularly, *δ* = 0 corresponds to no subtype signal in the controls and *δ* =1 corresponds to the same level of subtype signal in cases and controls. Presented is the performance in terms of absolute Pearson correlation under the different configurations, averaged over 20 simulations with vertical bars representing standard errors. The results are based on simulated data with N=1,400, *k*_0_ = 300, and 2,000 features.

**Figure S3:**
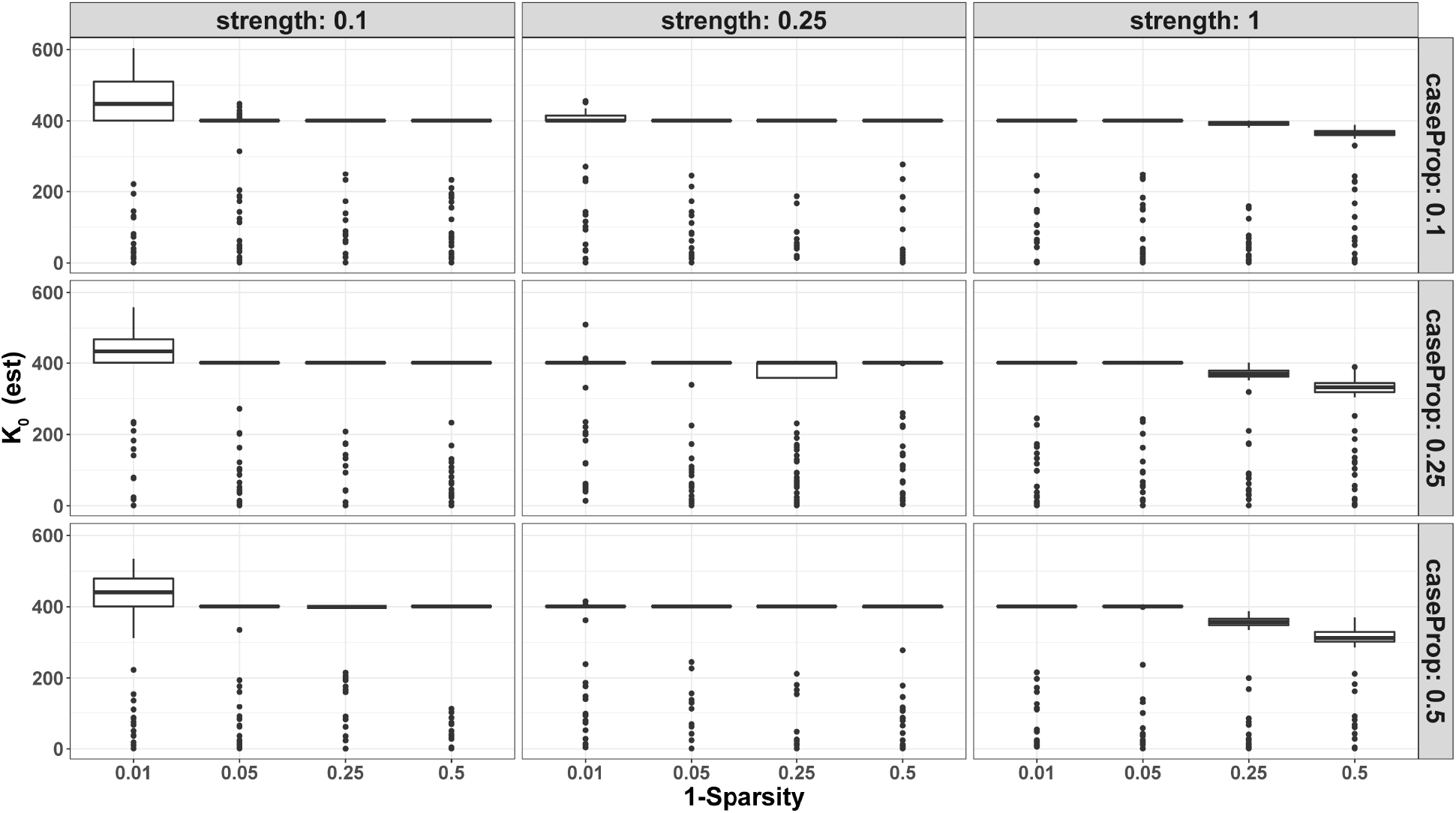
PACA estimates of the number of effective dimension of the shared variation between cases and controls (*k*) under simulations. The results are based on 100 simulations with *n* = 2000, *k*_0_ = 400 and *m* = 2000. The columns are evaluations at fixed signal strength, the rows are evaluations at fixed case proportions, and the X-axes indicate the level of signal sparsity (lower values mean more sparse).

**Figure S4:**
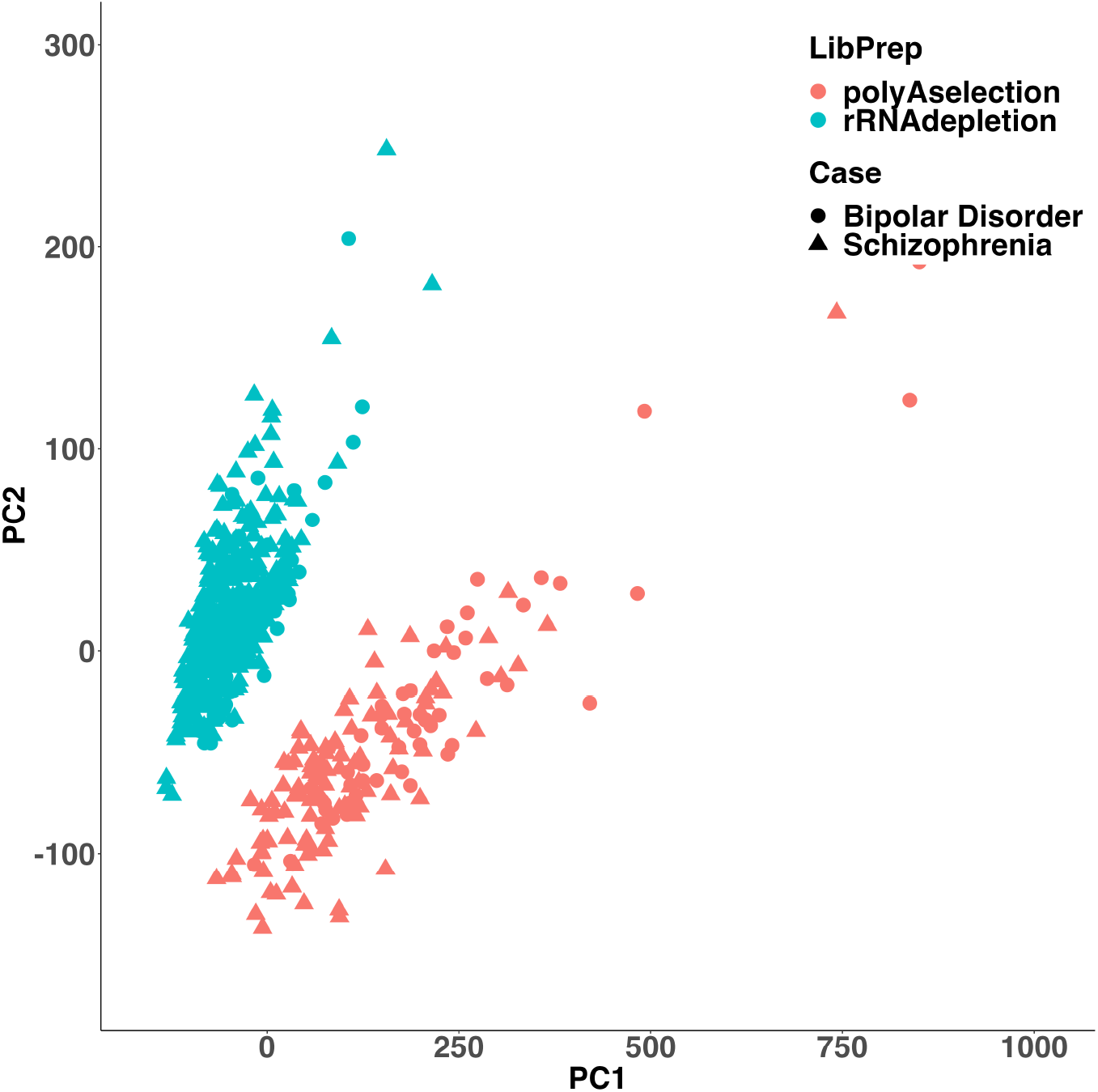
The top two principal components (PCs) of the pool of 472 schizophrenia cases and 172 bipolar disorder cases from the PsychENCODE data. The top two PCs perfectly separate the samples in the data based on the RNA Library preparation type, poly(A) selection (in blue) or rRNA depletion (in red).

**Figure S5:**
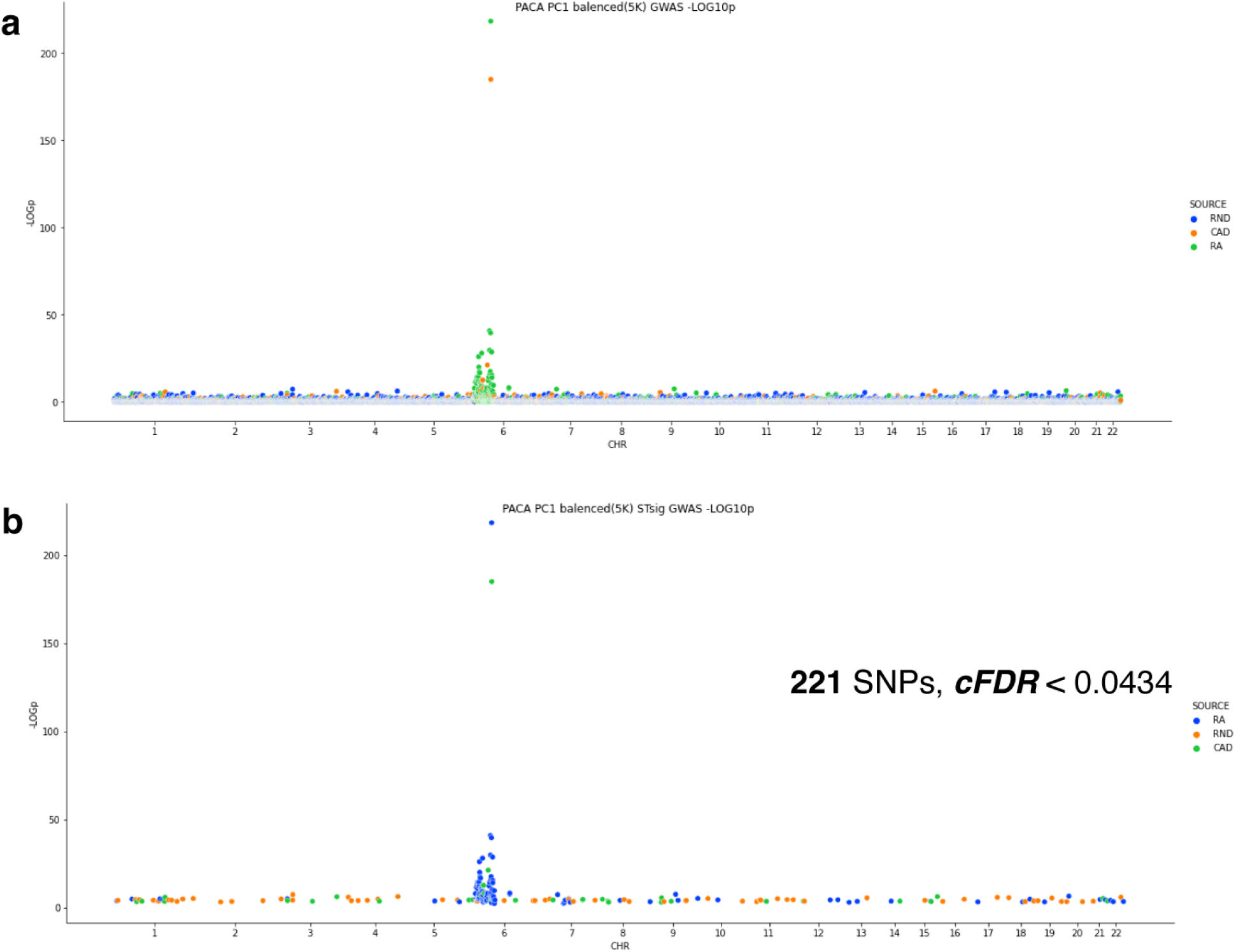
(a) Manhattan plot of a GWAS on pooled RA and CAD patients (N=10,000) with PACA PC1 as the phenotype. Results are shown for 10,000 SNPs, selected to be either RA-associated (green), CAD-associated (orange), or random (blue) SNPs. (b) Manhattan plot showing the 221 SNPs passing Subtest cFDR (<0.034). The 221 SNPs are either RA-associated (blue), CAD-associated (green), or random (orange) SNPs. The X-axes in the Manhattan plots correspond to chromosomal positions of the SNPs and the Y-axes reflect p-values on a -log_10_ scale.

### Part II

#### Supplementary Methods

##### S1 S1 Datasets

We used genotype data from the UK Biobank (UKBB) repsitory [1] and gene expression data from PsychENCODE [2, 3] for the null calibration experiment. We used both the aforementioned datasets and the genotype data of PsychENCODE for the validation analysis. In addition, we used the HapMap dataset [4] to illustrate null miscalibration.

##### S1.1 HapMap

We used genotypes from the HapMap project (release 23a, filtered) [4]. We considered a total of 120 samples (60 Yoruba in Ibadan, Nigeria (YRI) and 60 Northern/Western European Utah, U.S. (CEU) founders) and 1,872,641 SNPs for the analysis.

##### S1.2 Methylation Data

We used array DNA methylation datasets from the GALA II study with Latino population (n=573; GEO accession ID GSE77716) [5] and from the Hannum et al. study with European population (n=656; GEO accession ID GSE40279) [6]. We considered the top 50,000 most variable CpGs in the data. Prior to our analysis, we excluded outlier samples that demonstrated extreme values in the top two PCs of the data (values in PC1 or PC2 exceeding 3 standard errors), which resulted in n=558 and n=651 samples for the GALA II and Hannum datasets, respectively. Furthermore, the GALA II dataset includes 77 samples that are labeled as mixed Latino or other ethnicity; these were excluded from the final evaluation for which we only considered Mexican and Puerto Rican samples.

Mexican and Puerto Rican populations are known to be composed of European, Native American, and African populations. Therefore, the European samples in the Hannum data do not represent an ideal background dataset, given that information of European ancestry exists in the GALA II data. In our application of PACA we therefore employed a more relaxed assumption on the upper bound of the expected ratio between the variance of the subtype information in the Latino population compared with the variance of the subtype direction of information of the samples in the European population (δ = 4).

##### S1.3 UK BioBank

We used a curated set of 291,273 unrelated white British individuals (459,792 SNPs) from the UK BioBank data (application No. 33127). This version of the data was used to calculate the first 10 PCs, which were uses as covariates in all GWAS runs. We used Field 20002 (Non-cancer illness, self reported; instance 1) from the phenotypic data to obtain disease type and status, and we classified individuals as healthy controls if they had no reported diseases in Field 20002; eventually, we randomly selected a subset of 5,000 controls.

For our Coronary Artery Disease (CAD) and Rheumatoid Arthritis (RA) analysis we also used Field 20002 in order to find 2,500 mutually exclusive cases of CAD and RA (randomly sampled from the complete groups of cases). Our analysis defines CAD patients as any individual who reported Angina or myocardial infraction. We considered a set of 10,000 SNPs, including 2,000 RA-associated, 2,000 CAD-associated and 6,000 random (null) SNPs. The 2,000 RA-associated and 2,000 CAD-associated SNPs were selected as the top mutually exclusive 2,000 SNPs for each disease using an LD Clumped (± 100Kb window, LD < 0.1) list of the top SNPs from the Neal Lab summary statistics (v3; http://www.nealelab.is/uk-biobank/) present in our dataset. For the null SNPs, we randomly choose 6,000 SNPs from a LD pruned list (LD < 0.1) that had no overlap with the 2,011 and 9,696 most significant RA- and CAD-associated SNPs, respectively.

##### S1.4 PsychENCODE

We used genotypes, gene expression, and covariate information for 472 schizophrenia (SCZ), 172 bipolar disorder (BP), and 33 Alzheimer’s disease (AD) patients, as well as for 644 controls. For the validation analysis, we only used SCZ, BP and controls. We used all 5,312,508 SNPs and all samples to learn the top 10 PCs which we used a covariates in all GWAS runs. We standardized all the raw counts of the 25,774 genes in the gene expression data, and we used the a set of 217,863 LD pruned SNPs (< 0.1) for the Subtest analysis.

#### S2 Simulations

We generate complete simulations as outlined in this section for the power analysis. We simulated data with two subtypes using the following model. Let *X*_0_ ∈ ℝ^*m*×*n*_0_^, *X*_1_ ∈ ℝ^*m*×*n*_1_^ be the data for controls and cases, respectively, we assumed:

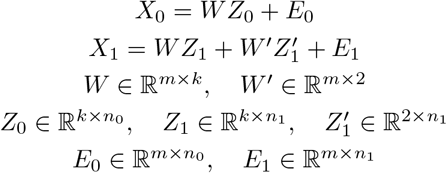

We simulated the the direction of the structure in the data, *W, W*’, as follows:

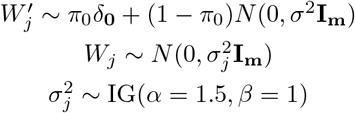

where *π*_1_ ∈ [0,1] reflects the signal sparsity. The signal sparsity controls the proportion of features with no effect (i.e., zero weights) in the *k* latent factors. 1 – can therefore be interpreted as the proportion of features affecting each of the *k* latent factors; for example, a fraction of 0.2 implies that a random combination of 20% of the m features define each of the *k* latent factors. We set *σ*^2^ = 3 in all of our simulations, and in the null case we set *W*’ = **0**.

Next, we simulate the structure in the shared and unique structure. Let *k* be the number of shared dimensions of structure across cases and controls. For *k*’ ≤ *k*, let *k*’ be the dim of the balanced shared variation, and *k*–*k*’ be the dim of the unbalanced shared variation. In the balanced axes of variation for *i* ∈ {1,…, *k*’}:

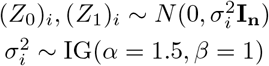

For the imbalanced axes of variation, when *k*’ ≠ *k, i* ∈ {*k*’ + 1,…, *k*}:

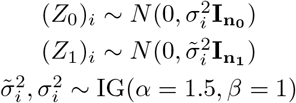

We simulate the subtype structure (membership), 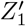, as one-hot encoded matrix assignment of individuals to the subtypes. We define “Prop” as the proportion of cases which belong to the first subtype. This is encoded by non-zero values in first column of 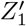. “Prop” helps us analyze the performance of methods under different levels of imbalance between the prevalence of subtypes. We can alter the subtype signal “Strength” by altering the 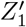 encodings’ non-zero values. “Strength” effectively scales the variance of the cases along the subspace which distinguishes the two subtypes. When defining a spectrum of phenotypic heterogeneity, instead of assigning individuals into subtypes, we sampled values from a standard random normal distribution.

We would also like to explore scenarios where the case specific subphenotypic signal is not exactly orthogonal to the structure in the controls. To explore this, we can also add some of the case specific structure to the controls 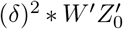. *W*’ is the same direction of structure as in the cases, 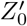 encode random assignment of controls to one of the two subgroups, and *δ* ∈ [0,1] controls the ratio of the variance of the subphenotypic signal in cases and the variance of the subphenotypic signal in controls. Pulling all this together we have the following model:

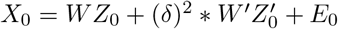

Finally, We simulate the noise terms from a normal distribution:

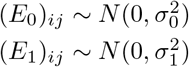

where 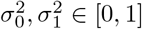 controls the strength of the noise added to the controls and cases, respectively.

#### S3 cPCA evaluation

The optimal choice of the contrastive hyperparameter *α* is unclear without prior information. We instead choose to perform an overly optimistic evaluation of cPCA in a setting where we have the subtype information and can estimate estimate the optimal contrast value. To do so for every run, we let cPCA automatically evaluate at 10 different contrastive hyperparameter values and only select the hyperparameter yielding the highest correlation with the subtype signal. This represents a setting where the subtype information is know*a priori* and could be considered the near best-case performance for cPCA. That being said, if we force cPCA to return a single contrastive hyperparameter it always, in our experiments, returns a value of 0. In such cases we use a small offset to set *α* = 0.01. As expect, the results seen at this setting were indistinguishable from PCA. In the PsychENCODE and methylation data analysis, we select the *α* which maximizes the correlation of cPC1 with the subtype of interest among the list of values suggested by cPCA. Finally, we also assess the results of cPCA at *α* = ∞ (INFcPCA). In practice its not possible to actually set the contrastive hyperparameter to ∞, instead we set it to an extremely high value (α = 10^6^).

#### S4 Evaluation Metrics

##### S4.1 Power

For performance evaluation, we use the Pearson’s correlation between a candidate subtype signal and the true subtype labels as a metric for power. To assess power, we use complete simulations instead of augmenting real data. There are several reasons we do this. Firstly, we want to assess methods under our model assumptions and its best to do this under complete simulations as we can control most properties of the data. Additionally, it is not trivial to add external structure to null genetic/gene expression data without fundamentally changing the nature of the data. Doing so could make it either trivial to detect the signal because it is distributionally distinct from the generative distribution of the base data or it could be stronger than any shared variation. Secondly, when using real data we do not have an *a priori* estimate of the number of shared dimensions between the case and controls, which makes it impossible to evaluate our *k*-selection procedure. Finally, our simulation framework allows us to control the presence/absence of imbalanced confounder to simulate more realistic data regimes and understand the impact of violating the orthogonality assumption. Such analysis is either not possible or quite hard when using real data since we have no control over the true undying shared structure.

##### S4.2 Null Calibration

We used a straight forward case/control label permutation approach to quantify the statistical significance of a tentative subtype signal at a given *k*, or *α* for cPCA. At a fixed *k*, we build a null distribution of variances of the top PC by running PACA on all the permuted datasets. Then use this empirical null distribution to quantify the statistical significance of the observed top PC’s variance. This procedure should be able to reject the null when there is sufficiently strong variation unique to the cases. The exact same procedure is also used to quantify the statistical significance of the top PCA and cPCA components.

Note that we do not run analysis for the Null case using simulations because we are able to evaluate under more realistic null settings when we randomly split (real) healthy samples randomly into cases and controls. The real data we use for null calibration evaluation may contain variation that stem from unknown and possibility imbalanced sources. These factors make this evaluation procedure a better proxy for the settings of real applications. We do acknowledge that the randomization of samples may not sufficiently suppress unknown variation that could be tagged as a possible subtype. This, however, simply means that our real data calibration results are conservative relative to evaluation with simulated data under a perfect/complete null, which we rarely observe in real data.

##### S4.3 Subtest

Finally, we use Subtest [7], to check if a particular axis of variation defines an axis of genetically different architecture. Subtest is a statistical test to determine the existence of phenotypic subgroups in genetics. It does so by modeling the distribution of all SNP association statistics and fits a mixture of Gaussians. Explicitly, it tries to fits SNPs to 3 Bivariate Gaussian: 1) a distribution of SNPs associated with neither the disease or proposed subtype, 2) a distribution of SNPs only significantly associated with the disease and not the subtype and 3) a distribution of SNPs which are either (i) associated only with the subtype or (ii) significantly associated with both the disease and proposed subtypes. Subtest fits Null model with a mixtures of model 1), 2) and 3)(i) and the alternate model with 1), 2) and 3)(ii), the difference of these model likelihoods provides a (pseudo) LogLikelihoods Ratio (pLR). We can generate a null distribution by permuting the subtype values, generating the null test statistics and fitting the Subtest to this data, which generates the null distribution for the pLR, which we can now use to derive a p-value quantify the the significance the proposed subtypes.

Fundamentally, Subtest assumes that if valid subtypes exist, they should have some non-negligible proportion of SNPs which are associated with the subtypes and the primary disease and it tries to find the existence of such subtypes. We don’t believe that this models encompasses the full space of all possible subtypes, nonetheless this a well-motivated and well-defined set space for validating putative subtypes. In our application, we are using genetics to validate proposed subtypes uncovered by PACA in gene expression or genetic data. Therefore, we can use Subtest to test whether PACA PC1 learns a valid axis of variation that defines a genetically differential subtype. Please also note that Subtest also provides a way to test for subtype differences in each SNP using a Bayesian conditional False Discovery Rate (cFDR), by conditioned the subgroup test statistic on the primary phenotype test statistic for each SNP. We can use this procedure to select for SNPs which tentatively discriminate the proposed subtypes. We only provide a very brief introduction to Subtest here, please refer to [7] for a more detailed explanation on the topic.

#### S5 Implementation of PACA

PACA is mainly implemented in R, however, for the core CCA worker function we used a recently developed highly efficient cpp implementation [8].

All analysis, execution and visualization were performed in a series of R and Python (for cPCA and PCA) scripts. PACA will be available as a R package which will be open sourced and make public upon publication.

#### S6 Randomized PACA

The CCA step in PACA imposes the limitation *m* ≥ max(*n*_0_, *n*_1_). In practice, there may be datasets in which this condition does not hold. In such cases, we can apply PACA on a subset of the samples, however, this is clearly sub-optimal to exclude data. In order to address this limitation, we introduce randomized PACA (rPACA), a randomized variant of PACA, which can operate under the setup *m* < max(*n*_0_, *n*_1_) and provide cases-specific variation for all *n*_1_ samples in *X*. rPACA is based on the idea that the full sample covariance matrix of the cases (after adjustment for shared sources of variation) can be approximated by learning projections from multiple subsets of the samples in the data. Finding the eigenvector that corresponds to the top eigenvalue of the estimated sample covariance matrix will then allow us to describe the most dominant variation in *X* while accounting for the sources of variation that are shared with *Y*. Below, we provide a brief description of the rPACA algorithm; see Supplementary Algorithm 2 for the complete details.

Briefly, rPACA begins by randomly splitting the data into two sets, each with a subset of the cases and a subset of the controls. rPACA uses one of the sets to learn the shared axes of variation between the cases and controls in the set, which are then used to adjust cases in both sets of the data. We then obtain the top *q* PCs of both sets of cases and stack them into a single matrix; of note, we match the variances of the projections across the two subsets of cases, along each of the *q* PCs, which alleviates possible variance shrinkage. Repeating this procedure multiple times using multiple splits of the data and combining the PCs across the multiple iterations allows us to use them to approximate the sample-covariance matrix in X. So we believe we should be able to randomly sub-sample the case/control data to approximate the full corrected case covariance matrix which can be used to find the top case variation specific PC.

##### Algorithm S1 Randomized PACA (rPACA)

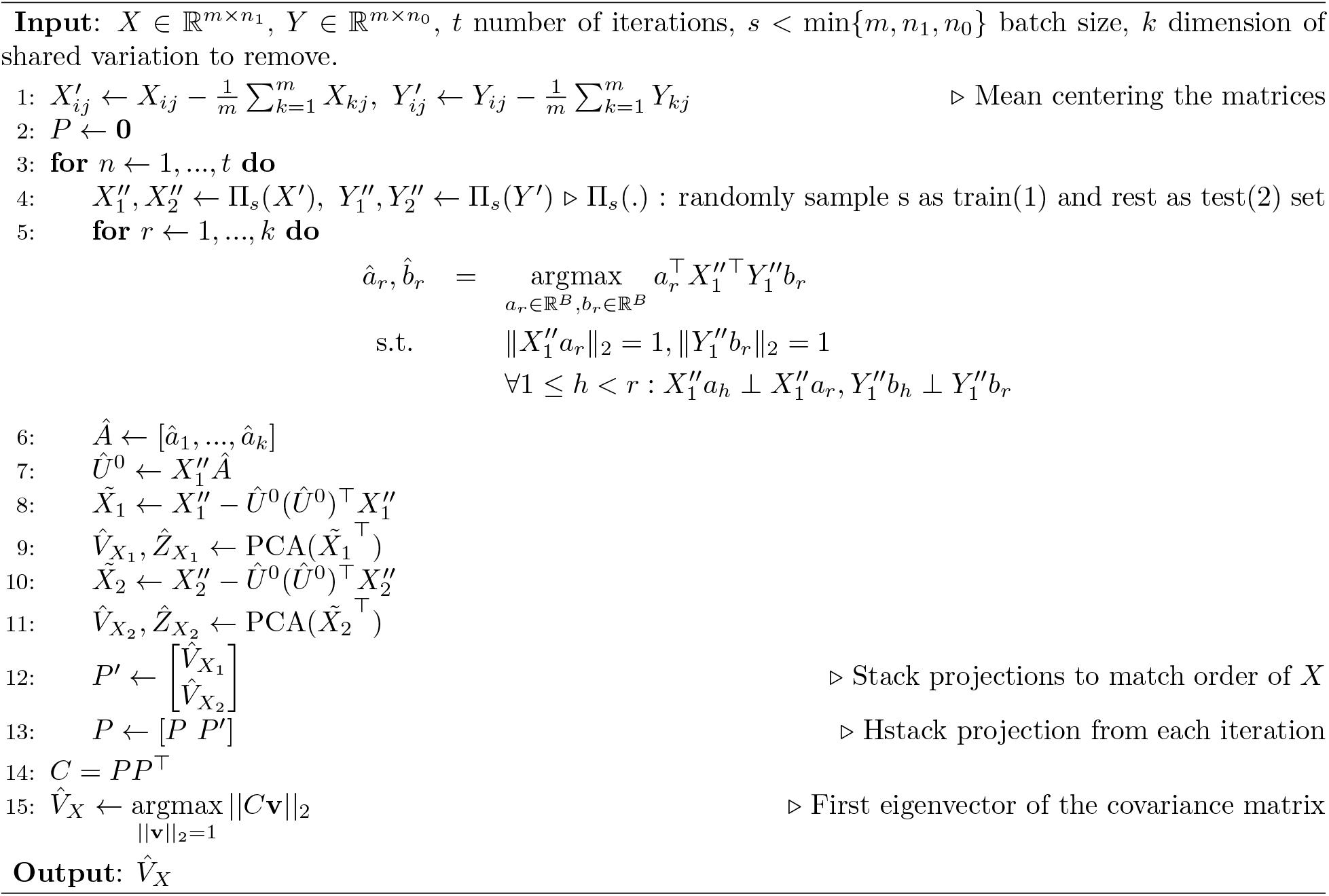

## References

[1] HP Himsworth. “Diabetes mellitus: Its differentiation into insulin-sensitive and insulin-insensitive types”. In: Diabetic Medicine 28.12 (2011), pp. 1440–1444.

[2] Juan-Sebasti’an Franco, Jenny Amaya-Amaya, and Juan-Manuel Anaya. “Thyroid disease and autoimmune diseases”. In: Autoimmunity: From Bench to Bedside [Internet]. El Rosario University Press, 2013.

[3] Subrata Manna and Marina K Holz. “Tamoxifen action in ER-negative breast cancer”. In: Signal transduction insights 5 (2016), STI–S29901.

[4] Jon McClellan and Mary-Claire King. “Genetic heterogeneity in human disease”. In: Cell 141.2 (2010), pp. 210–217.

[5] Annegien Broeks et al. “Low penetrance breast cancer susceptibility loci are associated with specific breast tumor subtypes: findings from the Breast Cancer Association Consortium”. In: Human molecular genetics 20.16 (2011), pp. 3289–3303.

[6] Montserrat Garcia-Closas et al. “Genome-wide association studies identify four ER negative–specific breast cancer risk loci”. In: Nature genetics 45.4 (2013), pp. 392–398.

[7] Shafali S Jeste and Daniel H Geschwind. “Disentangling the heterogeneity of autism spectrum disorder through genetic findings”. In: Nature Reviews Neurology 10.2 (2014), pp. 74–81.

[8] Vikas Bansal et al. “Genome-wide association study results for educational attainment aid in identifying genetic heterogeneity of schizophrenia”. In: Nature communications 9.1 (2018), pp. 1–12.

[9] Miriam S Udler et al. “Type 2 diabetes genetic loci informed by multi-trait associations point to disease mechanisms and subtypes: a soft clustering analysis”. In: PLoS medicine 15.9 (2018), e1002654.

[10] Andy Dahl and Noah Zaitlen. “Genetic influences on disease subtypes”. In: Annu. Rev. Genomics Hum. Genet 21 (2020), pp. 413–435.

[11] Luis B Barreiro et al. “Deciphering the genetic architecture of variation in the immune response to Mycobacterium tuberculosis infection”. In: Proceedings of the National Academy of Sciences 109.4 (2012), pp. 1204–1209.

[12] Benjamin P Fairfax et al. “Innate immune activity conditions the effect of regulatory variants upon monocyte gene expression”. In: Science 343.6175 (2014), p. 1246949.

[13] Mark N Lee et al. “Common genetic variants modulate pathogen-sensing responses in human dendritic cells”. In: Science 343.6175 (2014), p. 1246980.

[14] Alexander I Young, Fabian Wauthier, and Peter Donnelly. “Multiple novel gene-by-environment interactions modify the effect of FTO variants on body mass index”. In: Nature communications 7.1 (2016), pp. 1–12.

[15] Matthew R Robinson et al. “Genotype-covariate interaction effects and the heritability of adult body mass index”. In: Nature genetics 49.8 (2017), pp. 1174–1181.

[16] Alexander I Young, Fabian L Wauthier, and Peter Donnelly. “Identifying loci affecting trait variability and detecting interactions in genome-wide association studies”. In: Nature genetics 50.11 (2018), pp. 1608–1614.

[17] Rachel Moore et al. “A linear mixed-model approach to study multivariate gene–environment interactions”. In: Nature genetics 51.1 (2019), pp. 180–186.

[18] Andy Dahl et al. “A robust method uncovers significant context-specific heritability in diverse complex traits”. In: The American Journal of Human Genetics 106.1 (2020), pp. 71–91.

[19] Daniel Ferreira, Agneta Nordberg, and Eric Westman. “Biological subtypes of Alzheimer disease: A systematic review and meta-analysis”. In: Neurology 94.10 (2020), pp. 436–448.

[20] Joanne Ryan et al. “Phenotypic heterogeneity in dementia: a challenge for epidemiology and biomarker studies”. In: Frontiers in public health 6 (2018), p. 181.

[21] Judy H Cho and Peter K Gregersen. “Genomics and the multifactorial nature of human autoimmune disease”. In: New England Journal of Medicine 365.17 (2011), pp. 1612–1623.

[22] Jonathan Flint and Kenneth S Kendler. “The genetics of major depression”. In: Neuron 81.3 (2014), pp. 484–503.

[23] Roseann E Peterson et al. “Molecular genetic analysis subdivided by adversity exposure suggests etiologic heterogeneity in major depression”. In: American Journal of Psychiatry 175.6 (2018), pp. 545–554.

[24] Caroline M Nievergelt et al. “International meta-analysis of PTSD genome-wide association studies identifies sex-and ancestry-specific genetic risk loci”. In: Nature communications 10.1 (2019), pp. 1–16.

[25] Francesco Lescai and Claudio Franceschi. “The impact of phenocopy on the genetic analysis of complex traits”. In: PLoS One 5.7 (2010), e11876.

[26] Mirko Manchia et al. “The impact of phenotypic and genetic heterogeneity on results of genome wide association studies of complex diseases”. In: PloS one 8.10 (2013), e76295.

[27] Alexa A Woodward et al. “Genetic heterogeneity: Challenges, impacts, and methods through an associative lens”. In: Genetic Epidemiology (2022).

[28] Pooya Mobadersany et al. “Predicting cancer outcomes from histology and genomics using convolutional networks”. In: Proceedings of the National Academy of Sciences 115.13 (2018), E2970–E2979.

[29] Ming Tang et al. “The histologic phenotype of lung cancers is associated with transcriptomic features rather than genomic characteristics”. In: Nature communications 12.1 (2021), pp. 1–11.

[30] Charles M Perou et al. “Molecular portraits of human breast tumours”. In: nature 406.6797 (2000), p. 747.

[31] Dvir Netanely et al. “Expression and methylation patterns partition luminal-A breast tumors into distinct prognostic subgroups”. In: Breast Cancer Research 18.1 (2016), p. 74.

[32] Miriam Ragle Aure et al. “Integrative clustering reveals a novel split in the luminal A subtype of breast cancer with impact on outcome”. In: Breast Cancer Research 19.1 (2017), p. 44.

[33] Li Li et al. “Identification of type 2 diabetes subgroups through topological analysis of patient similarity”. In: Science translational medicine 7.311 (2015), 311ra174–311ra174.

[34] Martin Alda et al. “Investigating responders to lithium prophylaxis as a strategy for mapping susceptibility genes for bipolar disorder”. In: Progress in Neuro-Psychopharmacology and Biological Psychiatry 29.6 (2005), pp. 1038–1045.

[35] Fernando S Goes et al. “Mood-incongruent psychotic features in bipolar disorder: familial aggregation and suggestive linkage to 2p11-q14 and 13q21-33”. In: American Journal of Psychiatry 164.2 (2007), pp. 236–247.

[36] Jesse Mu et al. “Parkinson’s Disease Subtypes Identified from Cluster Analysis of Motor and Nonmotor Symptoms”. In: Frontiers in aging neuroscience 9 (2017), p. 301.

[37] Michael L Gatza et al. “A pathway-based classification of human breast cancer”. In: Proceedings of the National Academy of Sciences 107.15 (2010), pp. 6994–6999.

[38] Ash A Alizadeh et al. “Distinct types of diffuse large B-cell lymphoma identified by gene expression profiling”. In: Nature 403.6769 (2000), pp. 503–511.

[39] Therese Sørlie et al. “Gene expression patterns of breast carcinomas distinguish tumor subclasses with clinical implications”. In: Proceedings of the National Academy of Sciences 98.19 (2001), pp. 10869–10874.

[40] Huiyi Zhu et al. “Identification of 6 dermatomyositis subgroups using principal component analysisbased cluster analysis”. In: International journal of rheumatic diseases 22.8 (2019), pp. 1383–1392.

[41] Stacy L Sell et al. “Principal component analysis of blood microRNA datasets facilitates diagnosis of diverse diseases”. In: PloS one 15.6 (2020), e0234185.

[42] Smriti Prasad et al. “Principal components analysis as a tool to identify lesional skin patterns in cutaneous lupus erythematosus”. In: Journal of the American Academy of Dermatology 83.3 (2020), pp. 922–924.

[43] Md Rezaul Karim et al. “Deep learning-based clustering approaches for bioinformatics”. In: Briefings in Bioinformatics 22.1 (2021), pp. 393–415.

[44] Matan Hofree et al. “Network-based stratification of tumor mutations”. In: Nature Methods 10.11 (2013), 1108–1115.

[45] Marzieh Ayati, Mark R Chance, and Mehmet Koyutürk. “Co-phosphorylation networks reveal subtype-specific signaling modules in breast cancer”. In: Bioinformatics 37.2 (2020), 221–228.

[46] John Novembre et al. “Genes mirror geography within Europe”. In: Nature 456.7218 (2008), p. 98.

[47] Elior Rahmani et al. “Genome-wide methylation data mirror ancestry information”. In: Epigenetics & chromatin 10.1 (2017), pp. 1–12.

[48] Elior Rahmani et al. “Sparse PCA corrects for cell type heterogeneity in epigenome-wide association studies”. In: Nature methods 13.5 (2016), p. 443.

[49] Erika L Moen et al. “Genome-wide variation of cytosine modifications between European and African populations and the implications for complex traits”. In: Genetics 194.4 (2013), pp. 987–996.

[50] Richard T Barfield et al. “Accounting for population stratification in DNA methylation studies”. In: Genetic epidemiology 38.3 (2014), pp. 231–241.

[51] International HapMap Consortium et al. “The international HapMap project”. In: Nature 426.6968 (2003), p. 789.

[52] Lawrence A Loeb, Keith R Loeb, and Jon P Anderson. “Multiple mutations and cancer”. In: Proceedings of the National Academy of Sciences 100.3 (2003), pp. 776–781.

[53] Ahmet Zehir et al. “Mutational landscape of metastatic cancer revealed from prospective clinical sequencing of 10,000 patients”. In: Nature medicine 23.6 (2017), pp. 703–713.

[54] William S Bush and Jason H Moore. “Genome-wide association studies”. In: PLoS computational biology 8.12 (2012), e1002822.

[55] Yan Zhang et al. “Estimation of complex effect-size distributions using summary-level statistics from genome-wide association studies across 32 complex traits”. In: Nature genetics 50.9 (2018), p. 1318.

[56] Zhixiang Lin et al. “Simultaneous dimension reduction and adjustment for confounding variation”. In: Proceedings of the National Academy of Sciences 113.51 (2016), pp. 14662–14667.

[57] Abubakar Abid et al. “Exploring patterns enriched in a dataset with contrastive principal component analysis”. In: Nature communications 9.1 (2018), p. 2134.

[58] Abubakar Abid and James Zou. “Contrastive variational autoencoder enhances salient features”. In: arXiv preprint arXiv:1902.04601 (2019).

[59] Harold Hotelling. “Relations between two sets of variates”. In: Biometrika 28.3/4 (1936), pp. 321–377.

[60] KV Mardia, JT Kent, and JM Bibby. “Multivariate analysis, 1979”. In: Probability and mathematical statistics. Academic Press Inc (1979).

[61] Michael J. Gandal et al. “Transcriptome-wide isoform-level dysregulation in ASD, schizophrenia, and bipolar disorder”. In: Science 362.6420 (2018).

[62] Daifeng Wang et al. “Comprehensive functional genomic resource and integrative model for the human brain”. In: Science 362.6420 (2018).

[63] Joshua M Galanter et al. “Differential methylation between ethnic sub-groups reflects the effect of genetic ancestry and environmental exposures”. In: elife 6 (2017), e20532.

[64] Gregory Hannum et al. “Genome-wide methylation profiles reveal quantitative views of human aging rates”. In: Molecular cell 49.2 (2013), pp. 359–367.

[65] James Liley, John A Todd, and Chris Wallace. “A method for identifying genetic heterogeneity within phenotypically defined disease subgroups”. In: Nature genetics 49.2 (2017), pp. 310–316.

[66] Evan A. Boyle, Yang I. Li, and Jonathan K. Pritchard. “An expanded view of complex traits: From polygenic to omnigenic”. In: Cell 169.7 (2017), 1177–1186.

[67] Yan Zhang et al. “Estimation of complex effect-size distributions using summary-level statistics from genome-wide association studies across 32 complex traits”. In: Nature Genetics 50.9 (2018), 1318–1326.

[68] Clare Bycroft et al. “The UK Biobank Resource With Deep Phenotyping and genomic data”. In: Nature 562.7726 (2018), 203–209.

## References

[1] Clare Bycroft et al. “The UK Biobank Resource With Deep Phenotyping and genomic data”. In: Nature 562.7726 (2018), 203–209.

[2] Michael J. Gandal et al. “Transcriptome-wide isoform-level dysregulation in ASD, schizophrenia, and bipolar disorder”. In: Science 362.6420 (2018).

[3] Daifeng Wang et al. “Comprehensive functional genomic resource and integrative model for the human brain”. In: Science 362.6420 (2018).

[4] International HapMap Consortium et al. “The international HapMap project”. In: Nature 426.6968 (2003), p. 789.

[5] Elior Rahmani et al. “Sparse PCA corrects for cell type heterogeneity in epigenome-wide association studies”. In: Nature methods 13.5 (2016), p. 443.

[6] Gregory Hannum et al. “Genome-wide methylation profiles reveal quantitative views of human aging rates”. In: Molecular cell 49.2 (2013), pp. 359–367.

[7] James Liley, John A Todd, and Chris Wallace. “A method for identifying genetic heterogeneity within phenotypically defined disease subgroups”. In: Nature genetics 49.2 (2017), pp. 310–316.

[8] Mike Thompson et al. “Confined: Distinguishing biological from technical sources of variation by leveraging multiple methylation datasets”. In: Genome Biology 20.1 (2019).

